# Right lateralized posterior parietal theta-gamma coupling during sustained attention in mice

**DOI:** 10.1101/2020.06.11.147330

**Authors:** Ryan Yang, Corrinne Dunbar, Alina Sonesra, Suhyeorn Park, Timothy Pham, Rodney C. Samaco, Brett L. Foster, Atul Maheshwari

**Affiliations:** Department of Neurology, Baylor College of Medicine, Houston, TX; Rice University, Houston, TX; Department of Pharmacology, Baylor College of Medicine, Houston, TX; Department of Molecular and Human Genetics, Baylor College of Medicine, Houston, TX; Department of Neurosurgery, Baylor College of Medicine, Houston, TX; Department of Neuroscience, Baylor College of Medicine, Houston, TX

**Keywords:** cross-frequency coupling, electrocorticogram, gamma power, hemispheric asymmetry

## Abstract

Sustained attention is supported by circuits in the frontoparietal attention network. In human and primate studies, the right posterior parietal cortex (PPC) shows dominance for sustained attention, and phase-amplitude coupling (PAC) throughout the frontoparietal network correlates with performance on attention tasks. Here we evaluate oscillatory dynamics of bilateral PPC in mice during the 5-Choice Serial Reaction Time Task (5-CSRTT). Right PPC theta-gamma PAC (TG-PAC) and gamma power were independently elevated to a greater degree than the left PPC during the period prior to a correct response and were significantly correlated with accuracy in both simple and difficult tasks. Greater task difficulty was also associated with greater hemispheric asymmetry in TG-PAC, favoring the right PPC. These findings highlight the engagement of PPC with sustained attention in mice, reflected by increases in TG-PAC and gamma power, with maximal expression in the right hemisphere.

---

Attention is a critical process that relies heavily on integrating top-down and bottom-up cognitive processing for an organism to successfully achieve its behavioral goals (Desimone and Duncan, 1995). The ability to sustain attention to a task is mediated by the frontoparietal attention network (Desimone and Duncan, 1995; Sellers et al., 2016). Top-down processes from the medial prefrontal cortex (PFC) couple with bottom-up processes arising from primary sensory cortices to create a “salience map” in the posterior parietal cortex (PPC) (Arcizet et al., 2011). Determination of salience is necessary for planning and executing perceptually guided actions (Bucci, 2009; Corbetta and Shulman, 2002). Recent evidence in humans and non-human primates indicate that sustained attention is a rhythmic process reflected by dynamic changes in the electrocorticogram (ECoG) throughout the frontoparietal network (Fiebelkorn et al., 2018; Helfrich et al., 2018; Szczepanski et al., 2014). In addition, there is a consistent asymmetry of sustained attention favoring dominance of the right parietal lobe (Kim et al., 1999; Park et al., 2016). It has been argued that laterality of higher order functions such as language and attention has the evolutionary advantage of processing information efficiently by avoiding redundancy (Güntürkün et al., 2020). While work in rodents has shown that single neuron activity within PPC increases during detection of visual cues in a sustained attention task (Broussard et al., 2009), electrophysiological evidence of lateralized and/or rhythmic neural dynamics within PPC during sustained attention has been lacking.

Here we use an unbiased approach to evaluate dynamic frequency changes in bilateral PPC in mice during the 5-Choice Serial Reaction Time Task (5-CSRTT), a paradigm commonly utilized to study sustained attention in rodents (Carli et al., 1983; Chudasama and Robbins, 2004; Lustig et al., 2013; Robbins, 2002). In addition to changes in individual frequency bands, changes in phase-amplitude coupling (PAC) were also assessed. PAC is defined by the interaction between the phase of slower frequencies with the amplitude (or power) of faster frequencies (Canolty et al., 2006). Growing evidence points to the role of PAC in integrating signals across multiple spatiotemporal scales (Canolty and Knight, 2010). We dissect the roles of theta, gamma and theta-gamma phase-amplitude coupling (TG-PAC) during the 5-CSRTT with progressively increasing task difficulty. Independent of changes in theta and gamma power, we find a persistent dominance of right-hemispheric TG-PAC with increasing hemispheric asymmetry as task difficulty increases.

## Materials and Methods

### Animals

Adult male (> 8 weeks old) wild-type mice were maintained on a C57BL6/J background for over 10 generations from The Jackson Laboratory (Bar Harbor, Maine). Mice were housed in a vivarium with lights on between 8am to 8pm. Experiments were carried out according to the guidelines laid down by the Baylor Institutional Animal Care and Use Committee (IACUC).

### 5-Choice Serial Reach Time Task (5-CSRTT)

The 5-Choice Serial Reaction Time Task (5-CSRTT), an automated operant conditioning task that measures rodent visuo-spatial attention (Bari et al., 2008; Carli et al., 1983; Chudasama and Robbins, 2004; Robbins, 2002), was performed in a box with 5 apertures in a front curved wall and one aperture in the rear of the chamber (**Figure 1**, TSE Systems, Inc). Mice were food deprived to 85-90% of their free-feeding body weight as previously described (Bhandari et al., 2016) to ensure motivation for a food reward. Mice participated in attention testing 5 days per week (Monday-Friday) between 11am-3pm to prevent confounding diurnal variation. After 3 days of food deprivation, mice underwent a habituation phase for 10 days. During this phase, the mouse habituated to both the nose poke apertures and the association between correctly poking its nose in an illuminated aperture and the presentation of a pellet (Humby et al., 1999; Mar et al., 2013). Each trial began by poking the aperture at the rear of the chamber. One of the 5 front apertures was then pseudo-randomly illuminated. If the mouse correctly poked its nose into the illuminated aperture, it was rewarded with a 20 mg Dustless Precision Pellet (#F0071, Bio-Serv, Inc) from the same aperture. The target aperture remained illuminated until the mouse correctly poked its nose into that aperture. During this acclimatization phase, incorrect nose-pokes were not punished. After retrieving its reward, the mouse could again initiate a new trial at the back of the chamber. Each training session ended after 30 minutes. The mice then proceeded through a standard protocol (**Table 1**) as previously described (Bari et al., 2008).

**Table 1.**
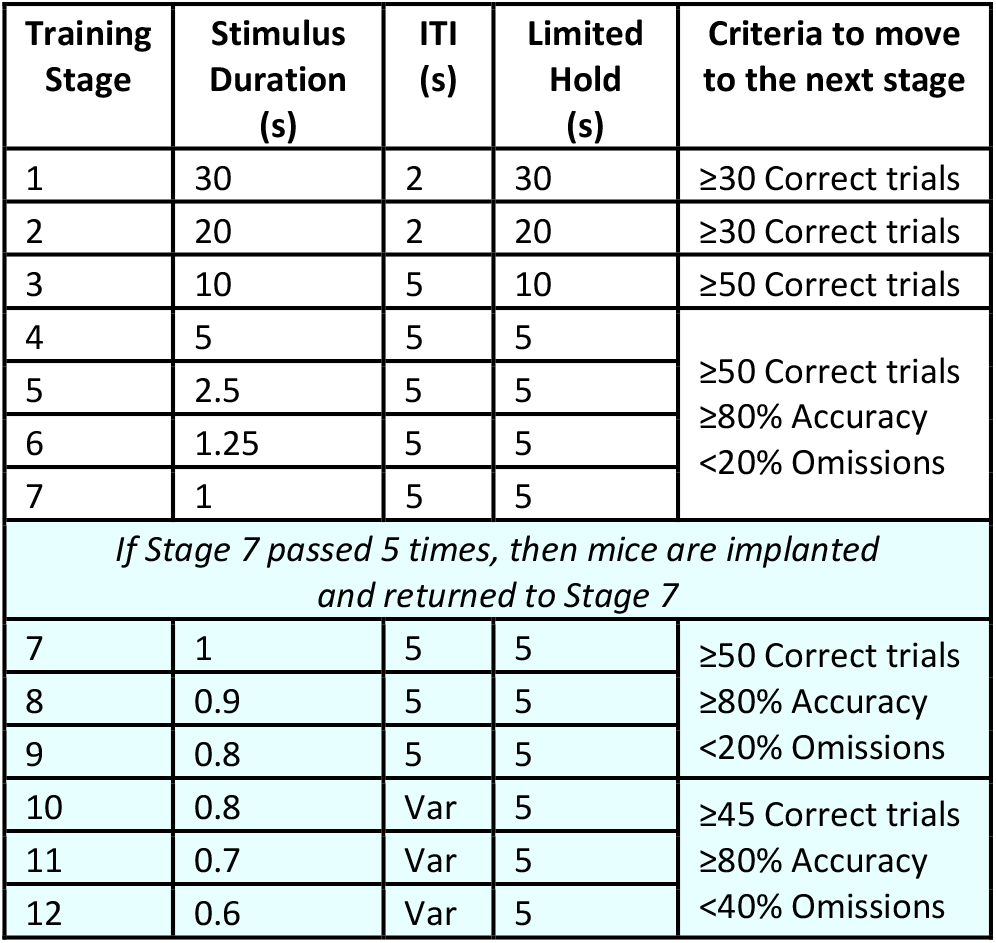
Protocol for Progression with 5-CSRTT. After habituation, mice progressed from Training Stage 1-7 with decreasing stimulus duration and limited hold periods, increasing inter-trial interval, and more stringent passing criteria (Bari et al., 2008). Var = variable.

After 10 days in the habituation phase, mice proceeded to the training phase. Starting at Training Stage 1, each session began with illumination of the rear aperture. The mouse initiated a trial and extinguished the illumination by poking its nose in the rear aperture. After the inter-trial interval (ITI), one of the 5 front apertures was pseudo-randomly illuminated. The mouse was able to poke its nose into the illuminated aperture and retrieve a reward any time during the stimulus duration (SD) or during a limited hold (LH) period, which was the period immediately following the SD (**Figure 1A-B**). If the mouse poked its nose into a different aperture (incorrect response), the chamber house light turned on to serve as a punishment. Omission responses, however, were not punished. After correct, incorrect or omission responses, the rear aperture was illuminated, and the mouse could then initiate another trial. The training protocol began with a SD of 30-s (SD30), an ITI of 2-s and a LH of 30-s, and became increasingly more difficult (with decreased SD and LH, and increased ITI) as the mice met the criteria to advance to the next stage (Bari et al., 2008; Bhandari et al., 2016). After successfully passing a stimulus duration of 1-s (SD1), mice were allowed to free-feed for 3 days in preparation for electrode implantation (see below). Post-operatively, the implanted mice were allowed to free-feed for an additional 2 days before they were placed back on the food deprivation protocol. Each implanted mouse proceeded with increasing task difficulty (shorter stimulus duration and introduction of a variable inter-trial interval of 2.5-5 seconds) until they reached a stimulus duration of 0.6 seconds with a variable inter-trial interval (SD0.6 vITI, **Table 1**). SD1 with a fixed ITI was defined as a simple task and SD0.6 vITI was defined as a difficult task based on prior studies (Fitzpatrick et al., 2018; Romberg et al., 2013). Mice were allowed to pass the SD0.6 vITI stage at least two times before completing the protocol.

**Figure 1.**
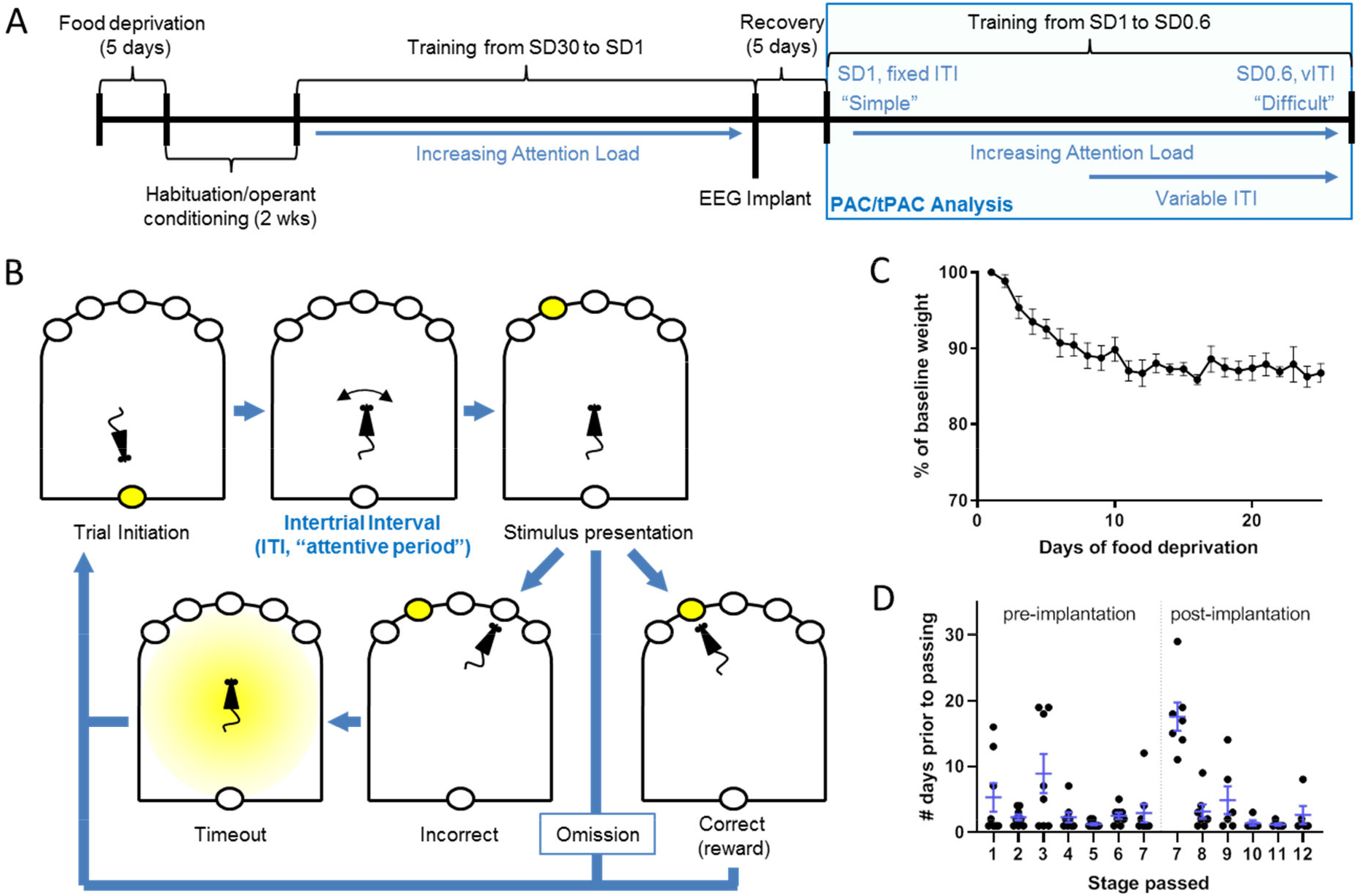
Schematic for 5-Choice Serial Reaction Time Task (5-CSRTT). (A) Prior to EEG implantation, mice were trained with increasing attention load until mastering a stimulus duration of 1 second (SD1). After EEG implantation, mice were returned back to SD1 and progressed with increasing attention load, including the addition of a variable inter-trial interval (vITI), until mastering SD0.6 with vITI. PAC/tPAC analysis was performed on the experiments within the blue box. (B) Mice initiated their own trials with the attentive period defined as the inter-trial interval, followed by stimulus presentation and one of 3 outcomes: correct response, incorrect response or omission. (C) Food deprivation caused consistent reduction in body weight. (D) Progression through the 5-CSRTT occurs with the greatest lag occurring at Stage 3 (first stage requiring 50 correct trials) and Stage 7 (the immediate post-implantation period).

### Surgical Electrode Implantation

During electrode implantation, mice were anesthetized with isoflurane (2-4% in O_2_) anesthesia and surgically and surgically implanted with silver wire electrodes (0.13 mm diameter) inserted into the epidural space over the posterior parietal cortex (PPC, 2.5 mm posterior and 1.5 mm lateral to bregma) (Lyamzin and Benucci, 2019; Zhang et al., 2016) and frontal cortex (supplementary motor cortex, 1 mm anterior and 1 mm lateral to bregma) as a control. Electrodes were placed bilaterally through cranial burr holes and attached to a micro-miniature connector (Omnetics, Inc) cemented to the skull. The reference and ground electrodes were placed over the left and right cerebellum, respectively. In one mouse, after completing the protocol, the reference and ground electrodes were swapped to ensure PAC was not affected by referencing the right versus left cerebellum. Access to food and water was available *ad libitum* for at least 2 days post-implantation prior to re-initiation of food deprivation. If the mice were in an extremely distressed or unwell state, they were euthanized via CO_2_ inhalation using an automated CO_2_ delivery system (SmartBox, Euthanex, Palmer, PA).

### Video-EEG data recording

Intracranial EEG and behavioral activity in freely moving mice were recorded using simultaneous video-EEG monitoring (AD Instruments, Inc). EEG signals were sampled at 2 kHz with a low-pass anti-alias filter at 1 kHz. Video capture was performed with a 640×480 pixel resolution at 30 frames/s.

### Video-EEG data pre-processing

Investigators were blinded to training stage prior to data analysis. EEG data was then screened for artifact under the supervision of a board-certified epileptologist (AM). Since the power of high frequency oscillations may be falsely measured when there are sharp electrographic contours (Kramer et al., 2008), identified artifacts were digitally extracted in EEGLab (Delorme and Makeig, 2004). Raw data was notch filtered in EEGLab with a 1 Hz window around 60 Hz, 120 Hz, and 180 Hz.

### Spectral Analysis

EEG power spectra were estimated using the spectral analysis function (pwelch.m) in EEGLab, which divides a time series into 8 equal segments with 50% overlap using a Hamming window (Delorme and Makeig, 2004). Absolute power (AP) was calculated for both left and right PPC leads, and relative power (RP) was calculated by dividing the AP for each frequency by the total power (TP; 2-200 Hz) (RP = AP/TP). RP was then normalized with a log transformation before comparison between mice (Jobert et al., 2013; Maheshwari et al., 2017). Statistical differences between groups at baseline were tested using a repeated measures 2-way ANOVA with Sidak post-tests comparing values at each frequency. Statistical significance was set at an adjusted *p*<0.05 at 2 or more consecutive frequencies to avoid spurious significance. All statistical analysis was performed using Prism 8, version 8.4.0, GraphPad, CA.

### Phase-Amplitude Coupling (PAC) Analysis

PAC analysis was performed using functions from EEGLab and Brainstorm (Tadel et al., 2011) in Matlab (Mathworks, Inc., Natick, MA, USA). The PAC algorithm within Brainstorm utilizes the mean vector length method of determining a ‘direct PAC’ measure (Özkurt and Schnitzler, 2011). Frequencies of 2-30 Hz (x-axis) for phase and 30-200 Hz (y-axis) for amplitude were evaluated as described previously (Maheshwari et al., 2017).

### Time-Resolved Power (tGamma, tTheta) and Phase-Amplitude Coupling (tPAC) Analysis

For each electrode, time-resolved power (tGamma, tTheta) was calculated in Brainstorm using a sliding window of 4 s with 0.4 s steps for analysis at SD1 (given an ITI of 5 seconds) and a sliding window of 2 s with 0.4 s steps for analysis at SD0.6 (given a vITI with a range of 2.5-5 s). Based on the identified maximal value from the PAC comodulogram for a given recording session, a 20 Hz window around the peak frequency for amplitude was used for tGamma analysis. Similarly, a 2 Hz window around the peak frequency for phase was used for tTheta analysis. A relative power time series for tGamma and tTheta was then calculated by dividing the absolute power at each time point by the median power during the recording session (given a non-Gaussian power distribution). Time-resolved PAC (tPAC) was calculated in Brainstorm in a similar manner (Samiee and Baillet, 2017), using the same sliding windows as tGamma and tTheta. The 2 Hz window around the peak frequency for phase and the 20 Hz window around the peak frequency for amplitude were averaged to generate absolute tPAC, and relative tPAC was calculated by dividing the absolute tPAC by the mean tPAC across the recording (given a Gaussian tPAC distribution). The attentive phase (ITI) was defined as the 4 or 2 s period (for SD1 and SD0.6, respectively) prior to stimulus onset. The reward phase was defined as the equivalent time period immediately after a correct response, when the mouse was consuming its reward. Average tPAC was defined as the mean tPAC across the recording session. Relative tGamma, tTheta, and tPAC during the attentive phase (ITI) prior to correct responses were then compared with the ITI prior to incorrect/omission responses, the reward phase, and either the average tPAC or median tGamma/tTheta. To determine hemispheric asymmetry, an asymmetry index was created by subtracting the change between the reward and correct periods for tTheta/tGamma/tPAC values in the right PPC from values in the left PPC. A positive asymmetry index favored the left PPC and a negative index favored the right PPC. To ensure effect size was not confounded by multiple replicates per mouse (Aarts et al., 2014), group analysis was performed with variation between mice treated as a random effect. When considering the differences between incorrect/omission periods and either the correct period or the reward period, a nested one-way ANOVA was used. When considering the differences between values of paired responses such as reward and correct periods or left and right hemispheres, a repeated measures 2-way ANOVA with mixed-effects modeling was used. Correction for multiple comparisons was performed, and the significance was set at an adjusted *p*<0.05. All statistical analysis was performed using Prism 8, version 8.4.0, GraphPad, CA.

## Results

A total of 34 wild-type C57 male mice initiated the attention protocol (**Figure 1A-B**), of which 9 passed the training stage with a stimulus duration of 1 second (SD1). Weight change with food deprivation and progression through the predefined stages occurred with little variability (**Figure 1C-D**). Of the mice that were implanted, 7 completed SD1, and 5 completed SD0.6 with a variable inter-trial interval (vITI) with at least 2 passing sessions.

### Right-lateralized dominance of TG-PAC during the 5-CSRTT

Spectral analysis between 2-200 Hz was performed over the entire 30 minute window for the first session passing the simple attention task (SD1, fixed ITI) and the first session passing the difficult attention task (SD0.6, variable ITI). For both the simple task (**Figure 2B**) and the difficult task (**Figure 2C**), there was a clearly defined peak above the 1/f curve at theta frequency (8-10 Hz). However, there was no difference at any frequency between left and right posterior parietal (PPC) electrodes, independent of task difficulty (F_1,784_=0.080, *p*=0.777 at SD1; F_1,784_=0.691, *p*=0.406 at SD0.6; repeated measures 2-way ANOVA with correction for multiple comparisons, *p*>0.05).

**Figure 2.**
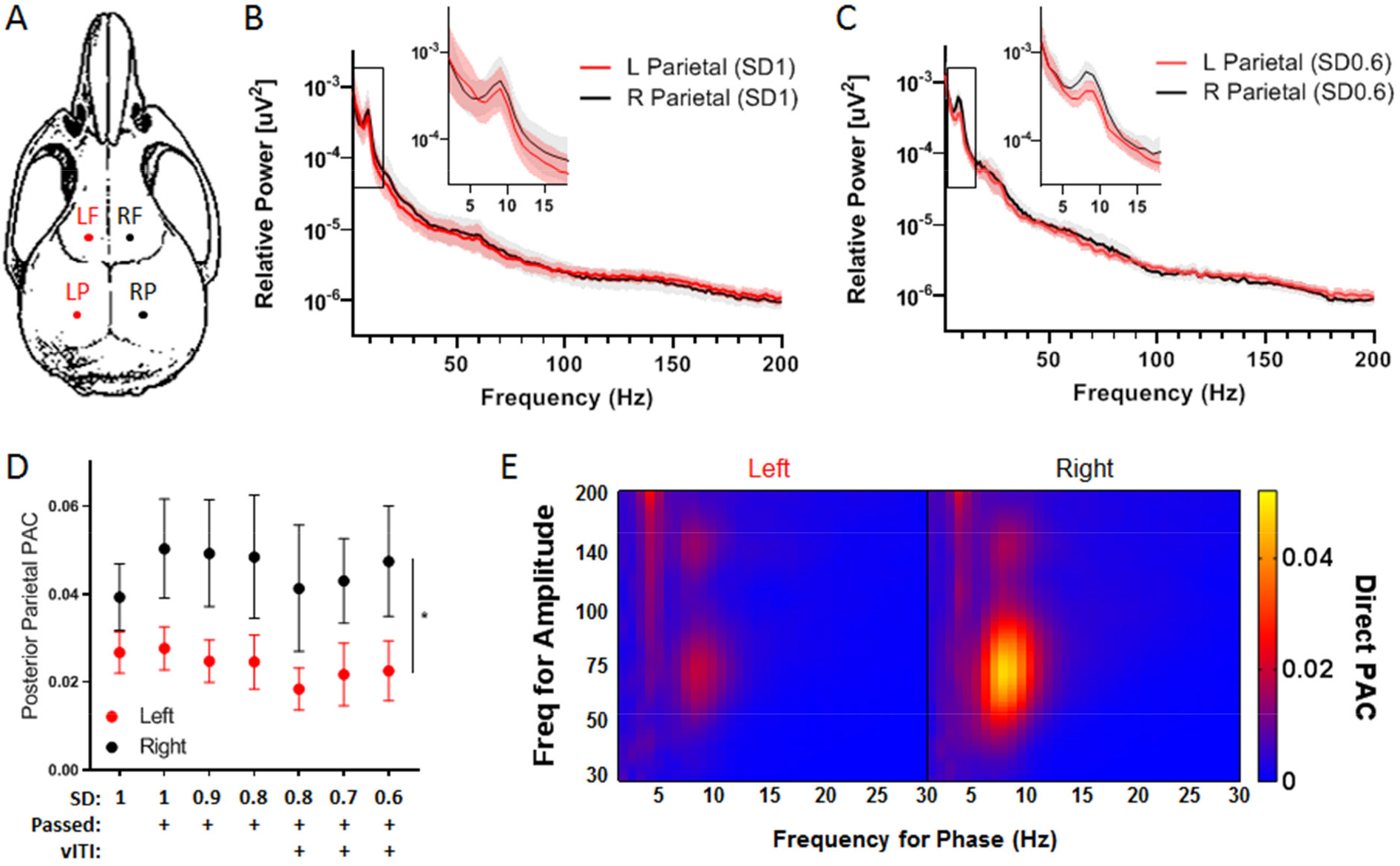
Lateralized Dominance of Posterior Parietal Theta-Gamma PAC (TG-PAC). (A) Coordinates for epidural electrode replacement, LF = left frontal, RF = right frontal, LP = left parietal, RP = right parietal (B) During SD1 (“simple task”), power spectra were equivalent over both hemispheres (n=7); inset showing theta peak. (C) Similarly, there were no differences overall during SD0.6 with vITI (n=5 mice, “difficult task”). (D) For the 5 mice that progressed to a stimulus duration of 0.6 seconds with a variable inter-trial interval (SD0.6, vITI), TG-PAC remained consistently elevated over the right PPC (F_1,4_=12.02, \**p*=0.026, repeated measures 2-way ANOVA). (E) Average comodulogram for left vs right PPC during SD0.6 vITI.

PAC was then analyzed during the overall recording sessions to evaluate differences between hemispheres with progressively increasing task difficulty. Baseline theta-gamma phase-amplitude coupling (TG-PAC) was identified in both left and right PPC during the first recording performed 1 week post implantation at SD1 (**Figure 2D**). As training progressed from SD1 to SD0.6 with vITI, TG-PAC in the right PPC was consistently elevated compared to the left PPC (F_1,4_=12.02, *p*=0.026, **Figure 2D**). The average comodulogram showed a distinct island of theta-gamma coupling that was greater in the right PPC when compared to the left PPC (maximal at 8.5 Hz for phase, and 73 Hz for amplitude, **Figure 2E**). Therefore, TG-PAC overall demonstrated a right-sided dominance during an attention task, despite no significant difference in power between hemispheres at any frequency.

### Right posterior parietal theta-gamma coupling correlates with accuracy during a simple attention task

To evaluate whether TG-PAC was specifically related to accuracy with task-based attention, time-resolved phase-amplitude coupling (tPAC) was subsequently analyzed, starting with sessions where mice passed SD1 (simple task). There was a consistent increase in right PPC tPAC values from the reward period to the highly attentive phase of the inter-trial interval period before a correct response (**Figure 3A**). In aggregate, tPAC in the right PPC during the ITI prior to a correct response significantly increased when compared to average (n=7, *p*=0.011, repeated measures one-way ANOVA with Sidak’s correction for multiple comparisons, **Figure 3B**). When compared to the reward period, relative tPAC was significantly increased in right PPC regions but not in bilateral frontal regions or left PPC (n=7, *p*=0.002, **Figure 3C**).

**Figure 3.**
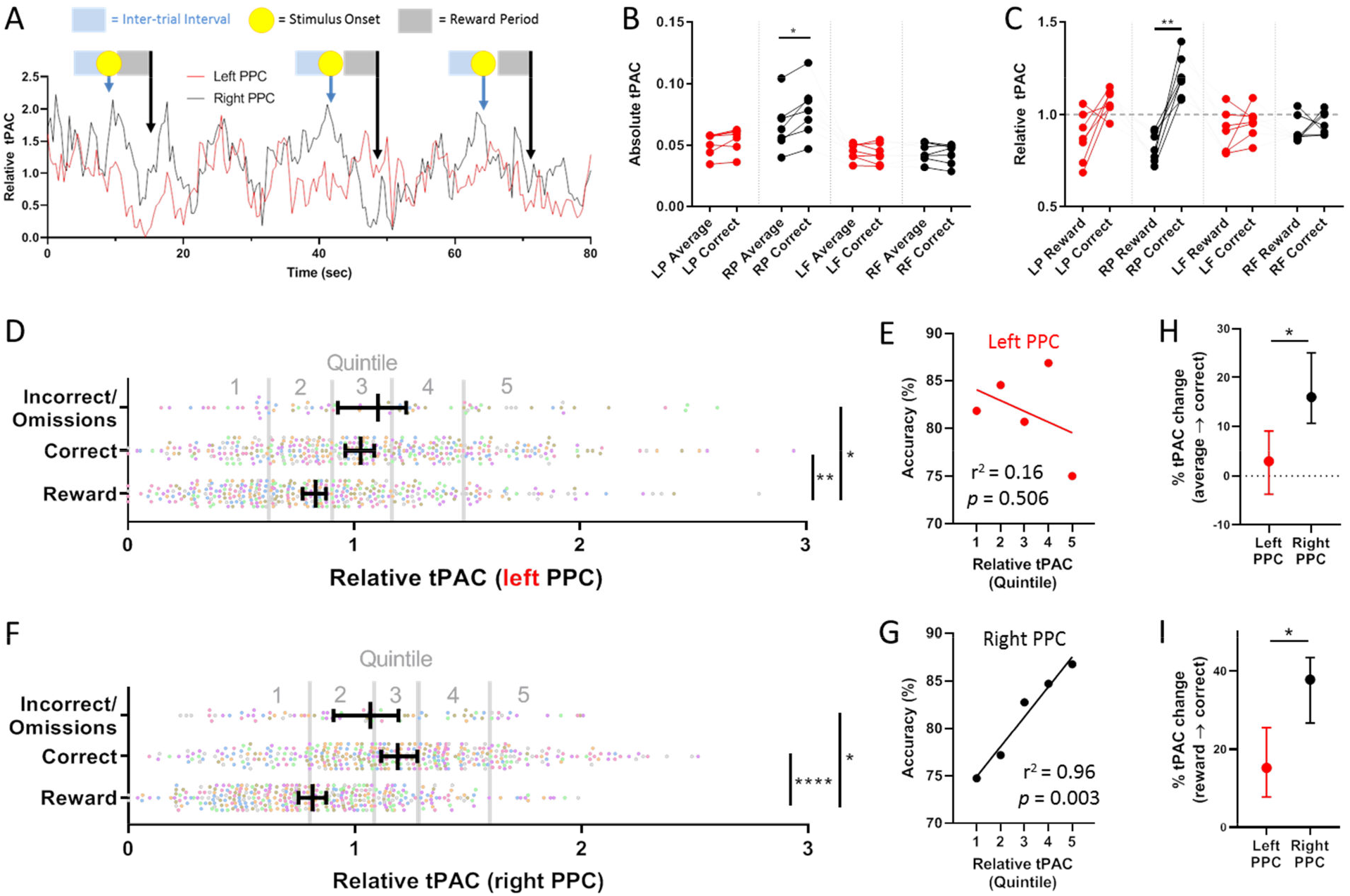
Positive correlation between accuracy and time-resolved PAC (tPAC) in the right posterior parietal region in a simple task (SD1). (A) Representative example of right PPC tPAC increasing during the inter-trial interval prior to a correct response, and then falling during the reward period with a stimulus duration of 1 second (using a 4-second sliding window). (B) Compared to average tPAC, only the right PPC had a significant increase in tPAC during the inter-trial interval prior to a correct response (\**p=*0.011, repeated measures one-way ANOVA, n=6 at SD1; LP = left parietal, RP = right parietal, LF = left frontal, RF = right frontal). (C) Compared to the reward period, both only the right PPC tPAC significantly increased in the interval prior to a correct response (\*\**p*=0.002, repeated measures one-way ANOVA). (D) With aggregate data from all mice at SD1 (n=7), there was a significant increase in tPAC from the reward period to the ITI before correct responses as well as the ITI before incorrect/omission responses in the left PPC (\**p*=0.016, \*\**p*=0.002, nested ANOVA, mixed-effects modeling). Each color represents a different mouse. (E) No significant relationship between relative tPAC prior to a correct response (binned by quintile) and accuracy in the 5-CSRTT in the left PPC. (F, G) In the right PPC, similar plots revealed significantly increased relative tPAC prior to correct and incorrect/omission responses (\**p*=0.012, \*\*\*\**p*<0.0001), and there was a significant correlation between tPAC and accuracy. (H) Significantly greater increase in tPAC prior to a correct response in the right compared to the left PPC when compared to average tPAC (\**p*=0.017, repeated measures 2-way ANOVA, mixed-effects modeling), or (I) when compared to the reward period (\**p*=0.015).

In the simple task (SD1), left PPC tPAC was significantly increased from the reward to the ITI prior to correct responses in 4 out of 7 mice, while right PPC tPAC was significantly increased in all 7 mice (*p*<0.05, Wilcoxon matched-pairs signed rank test, **Supplemental Figure 1**). To ensure the reference in the left cerebellum did not alter changes in theta-gamma coupling, PPC recordings were referenced to the right cerebellum in one mouse after referencing to the left cerebellum on the previous day, and the changes in tPAC were unchanged (**Supplemental Figure 2**). Since each session only had between 6-17 incorrect and omission responses, the ability to evaluate the relationship between accuracy and tPAC for individual mice was limited. Therefore, trials from all mice that passed SD1 (simple task) were aggregated, and changes in tPAC between the reward period, the ITI prior to correct responses, and the ITI prior to incorrect/omission responses were evaluated. Relative tPAC was significantly increased from the reward period to the ITI in both correct and incorrect/omission responses, but there was no significant difference between correct and incorrect/omission responses in either left or right PPC (**Figure 3D, F**, respectively). To further dissect the relationship between tPAC and accuracy, the tPAC for correct trials was binned into 5 equivalent quintiles, and accuracy (correct trials/total trials) was plotted for each quintile. The resultant vector showed no correlation in the left PPC (r^2^=0.16, *p*=0.506, **Figure 3E**); in contrast, there was a strong and significant positive correlation between tPAC and accuracy in the right PPC (r^2^=0.96, *p*=0.003, **Figure 3G**). In addition, directly comparing the ITI prior to correct responses in the left and right PPC, tPAC increased 3.0% (−3.7 to 9.1, 95% CI) above the average tPAC in the left PPC compared to 16.0% (10.7-25.1, 95% CI) in the right PPC (*p*=0.017, repeated measures 2-way ANOVA, mixed-effects modeling, **Figure 3G**). Compared to the reward period, left PPC tPAC was increased by 15.2% (7.8-25.6, 95% CI) in the ITI prior to correct responses, while the right PPC increased by 37.9% (26.7-43.5, 95% CI, *p*=0.015, **Figure 3H**). Therefore, while tPAC was elevated in bilateral PPC during the inter-trial interval, the right PPC was elevated to a greater degree and exclusively demonstrated a strong relationship with accuracy in a simple task of attention.

### Right posterior parietal gamma power independently correlates with accuracy during a simple task

Since tPAC is determined by the covariation in the amplitude of gamma power with the phase of theta rhythms, the relationship between time-resolved gamma power (tGamma) and attention-related performance was examined further. In the left PPC during a simple attention task, tGamma increased significantly from the reward phase to the ITI before correct responses (*p*<0.0001), but there was no significant relationship between left PPC tGamma and accuracy (r^2^=0.10, *p*=0.612, **Figure 4A-B**). In the right PPC, tGamma was significantly elevated during the inter-trial intervals prior to both correct and incorrect/omission responses compared to the reward period (p<0.0001 and *p*=0.017, respectively, **Figure 4C**). Similar to tPAC for the right PPC, there was a significant positive correlation between tGamma (prior to correct responses, by quintile) and accuracy (r^2^=0.89, *p*=0.017, **Figure 4D**).

**Figure 4.**
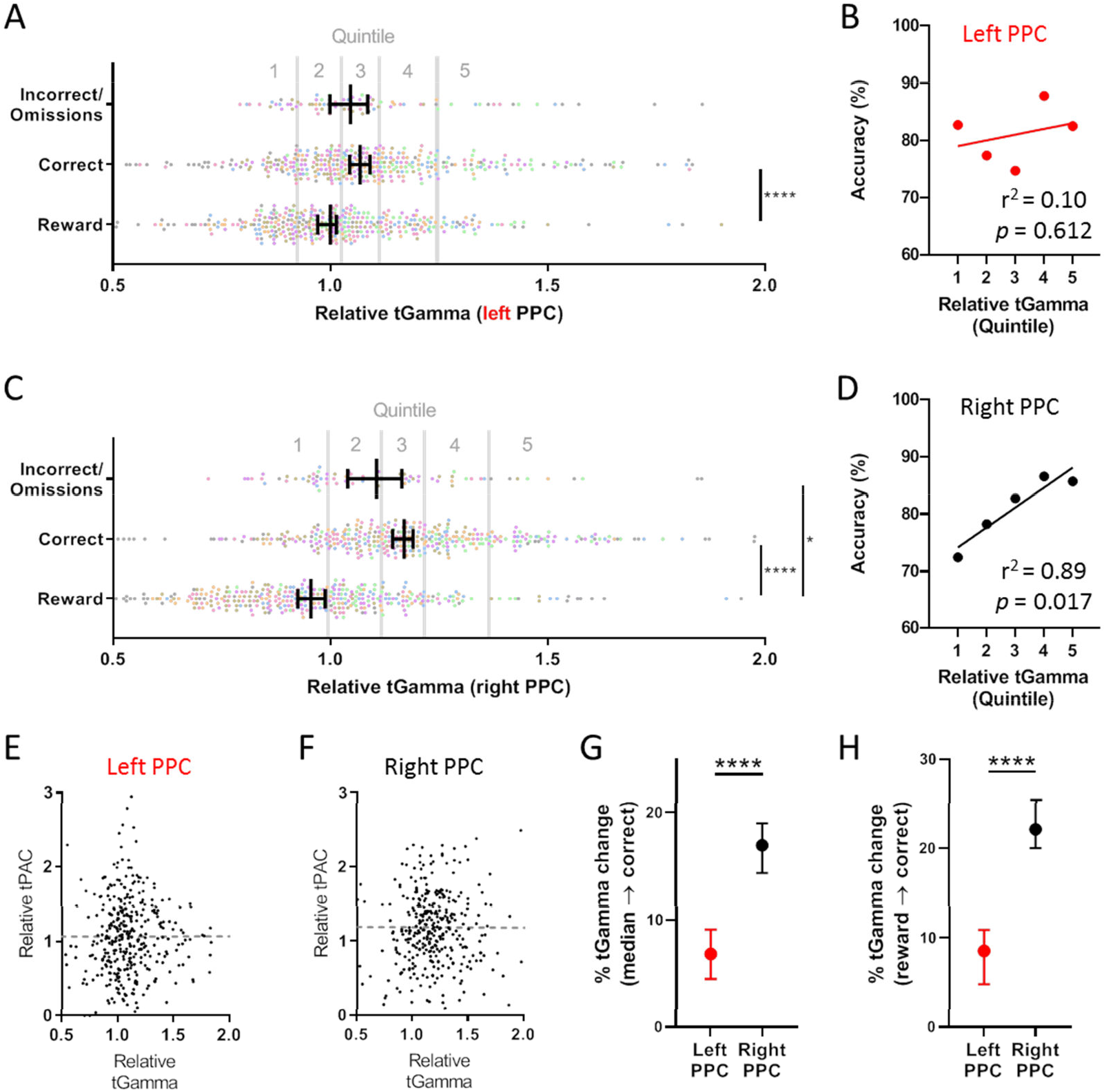
Relationship between time-resolved gamma power (tGamma) and performance at SD1 (simple task). (A) In the left PPC, there was a significant increase in tPAC from the reward period to the ITI before correct responses (\**p*<0.0001), with (B) no significant relationship between tGamma and accuracy. (C) In contrast, in the right PPC, gamma power was significantly increased significantly prior to both correct and incorrect/omission responses compared to the reward period (\**p*=0.017, \*\*\*\**p=*0.001). (D) Significant positive correlation between right PPC tGamma and accuracy. (E) No significant relationship between tGamma and tPAC in the left PPC (r^2^<0.0001, *p*=0.986) or (F) the right PPC (r^2^<0.0001; *p*=0.940). The magnitude of tGamma change prior to correct responses was significantly greater in the right PPC compared to the left PPC when referenced to either (G) median tGamma or (H) the reward period (\*\*\*\**p*<0.0001).

Importantly, there was no significant relationship between tGamma and tPAC prior to correct trials in either the left or right PPC (r^2^<0.0001, *p*=0.986 and r^2^<0.0001, *p*=0.940, respectively, **Figure 4E, F**). Directly comparing left and right PPC prior to correct responses, tGamma increased 6.8% (4.5-9.1, 95% CI) above median tGamma in the left PPC prior to correct responses and 16.9% (14.4-19.0, 95% CI) above median tGamma in the right PPC (*p*<0.0001, **Figure 4G**). Compared to the reward period, tGamma increased 8.5% (4.8 to 10.9, 95% CI) in the left PPC and 22.1% (20.0-25.4, 95% CI) in the right PPC (*p*<0.0001, **Figure 4H**). These findings indicate that right PPC gamma power has a greater association with attention performance than left PPC gamma power, independent of its contribution to tPAC in a simple task; hence, elevated tPAC is not simply driven by elevated gamma power.

### Right posterior parietal theta-gamma coupling correlates with accuracy during a difficult attention task

To determine how increasing task difficulty alters oscillatory dynamics during an attention task, 5 mice successfully progressed and completed 2 sessions with a stimulus duration of 0.6 seconds and a variable inter-trial interval (SD0.6 vITI, “difficult task”), each with at least 45 correct trials. Aggregating all data from these 10 sessions, left PPC tPAC was modestly but significantly increased from the reward period to the ITI prior to correct responses (*p*=0.020), but similar to the simple task, there was no significant relationship between accuracy and left PPC tPAC by quintile (**Figure 5A-B**). Right PPC tPAC, however, was significantly increased during the ITI prior to all responses, with an overall significantly greater increase before correct responses compared to incorrect/omission responses (**Figure 5C**). As in SD1, a significant positive correlation between tPAC and accuracy was observed (**Figure 5D**). Compared to average tPAC, the left PPC tPAC decreased by 0.6% (−9.2 to +4.5, 95% CI) while the right PPC tPAC increased by 27.2% (20.6-32.2, 95% CI) prior to a correct response (*p*<0.0001, repeated measures 2-way ANOVA, mixed-effects modeling, **Figure 5E**). Compared to tPAC during the reward period, the left PPC tPAC increased by 8.6% (1.4 to 13.8, 95% CI) while the right PPC tPAC was elevated to a significantly greater degree of 51.6% (45.7 to 57.5, 95% CI, **Figure 5F**). These findings support a dominance of right PPC TG-PAC that is maintained in both a simple and difficult task.

**Figure 5.**
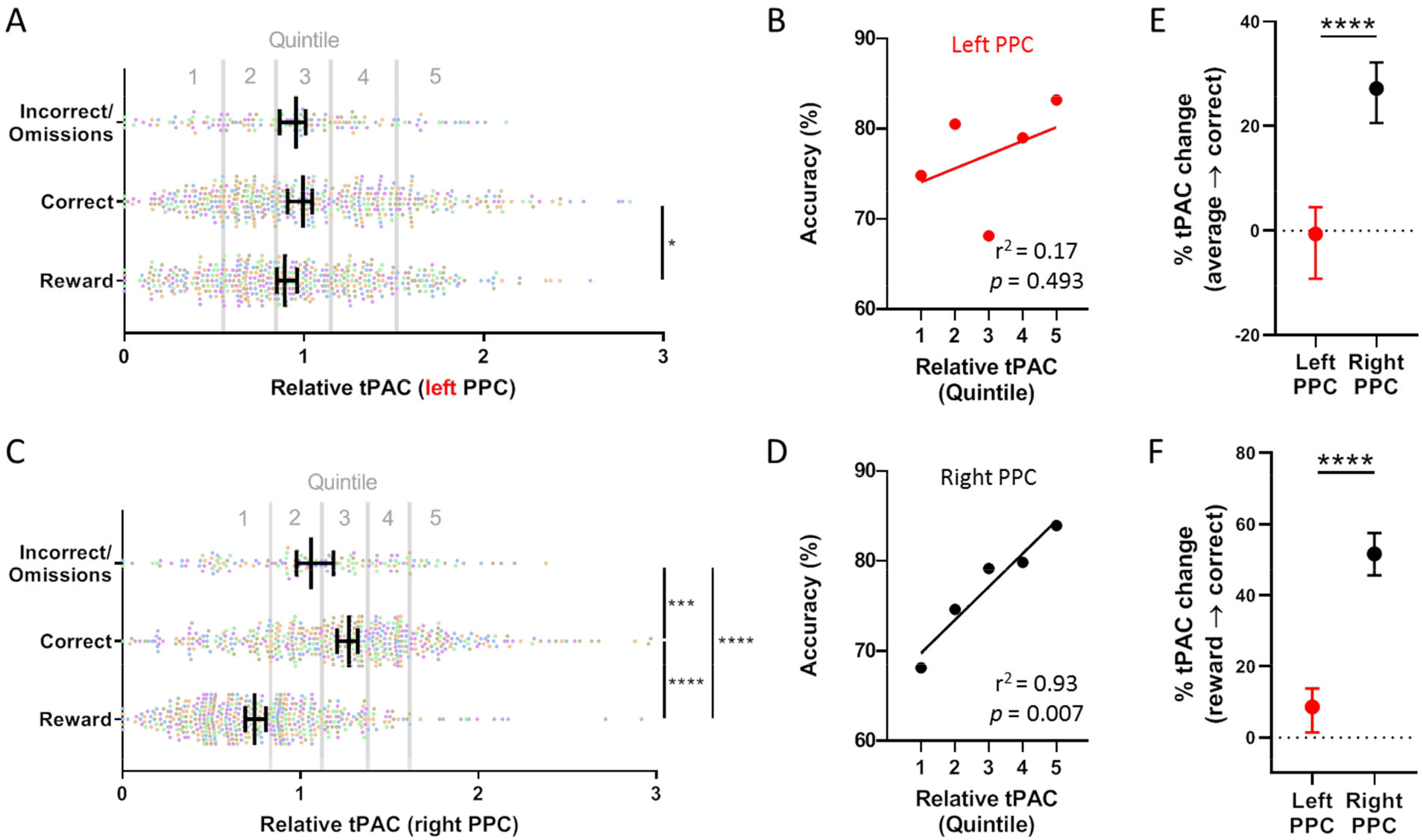
Dominant right-hemispheric tPAC at SD0.6 with vITI (difficult task). (A) In the left PPC, there was a significant increase in tPAC between the reward and correct response periods (n=5 mice, 2 sessions/mouse, \**p*=0.020), and (B) no significant relationship between tPAC and accuracy. (C) However, in the right PPC, there was significantly increased tPAC in the ITI before correct and incorrect/omission responses compared to the reward period as well as significantly increased tPAC with correct compared to incorrect/omission responses (\*\*\**p*<0.001, \*\*\*\**p*<0.0001). (D) Strong positive linear correlation between tPAC and accuracy in the right PPC. (E) Compared to average tPAC and (F) tPAC during the reward period, the right PPC increased to a significantly greater degree than the left PPC (\*\*\*\**p*<0.0001).

### Bilateral posterior parietal gamma power correlates with accuracy during a difficult task

Gamma frequency dynamics during a difficult task (SD0.6 vITI) were evaluated next. Left PPC tGamma was increased significantly in the ITI prior to correct responses compared to the reward period (*p*<0.0001, **Figure 6A**). In contrast to both tGamma in SD1 and tPAC in SD0.6, there was a significant positive correlation between accuracy and tGamma quintile in the left PPC (**Figure 6B**). Similar to the left PPC, right PPC tGamma was also significantly elevated in the ITI prior to correct responses compared to the reward period (*p*<0.0001, **Figure 6C**), and there was a significant positive correlation between accuracy and right PPC tGamma prior to correct responses when divided into quintiles (**Figure 6D**). There remained near-zero correlation between tGamma and tPAC in both left and right PPC (r^2^<0.0001, *p*=0.920 and r^2^=0.009, *p*=0.040, respectively, **Figure 6E, F**). Relative to median tGamma, left PPC tGamma increased by 16.3% (13.3-18.1, 95% CI) whereas right PPC tGamma increased to a significantly greater degree by 27.7% (24.4-30.7, 95% CI) prior to correct responses (*p*<0.0001, repeated measures 2-way ANOVA, mixed-effects modeling, **Figure 6G**). Compared to the reward period, left PPC tGamma increased by 10.1% (8.4-12.3, 95% CI) while right PPC tGamma increased to a significantly greater degree by 28.3% (26.0-30.0, 95% CI, *p*<0.0001, **Figure 6H**). These findings indicate that gamma power in the right PPC increases to a greater degree than the left PPC with a difficult task, but in contrast to a simple task, left PPC gamma power had a strong and significant correlation with accuracy.

**Figure 6.**
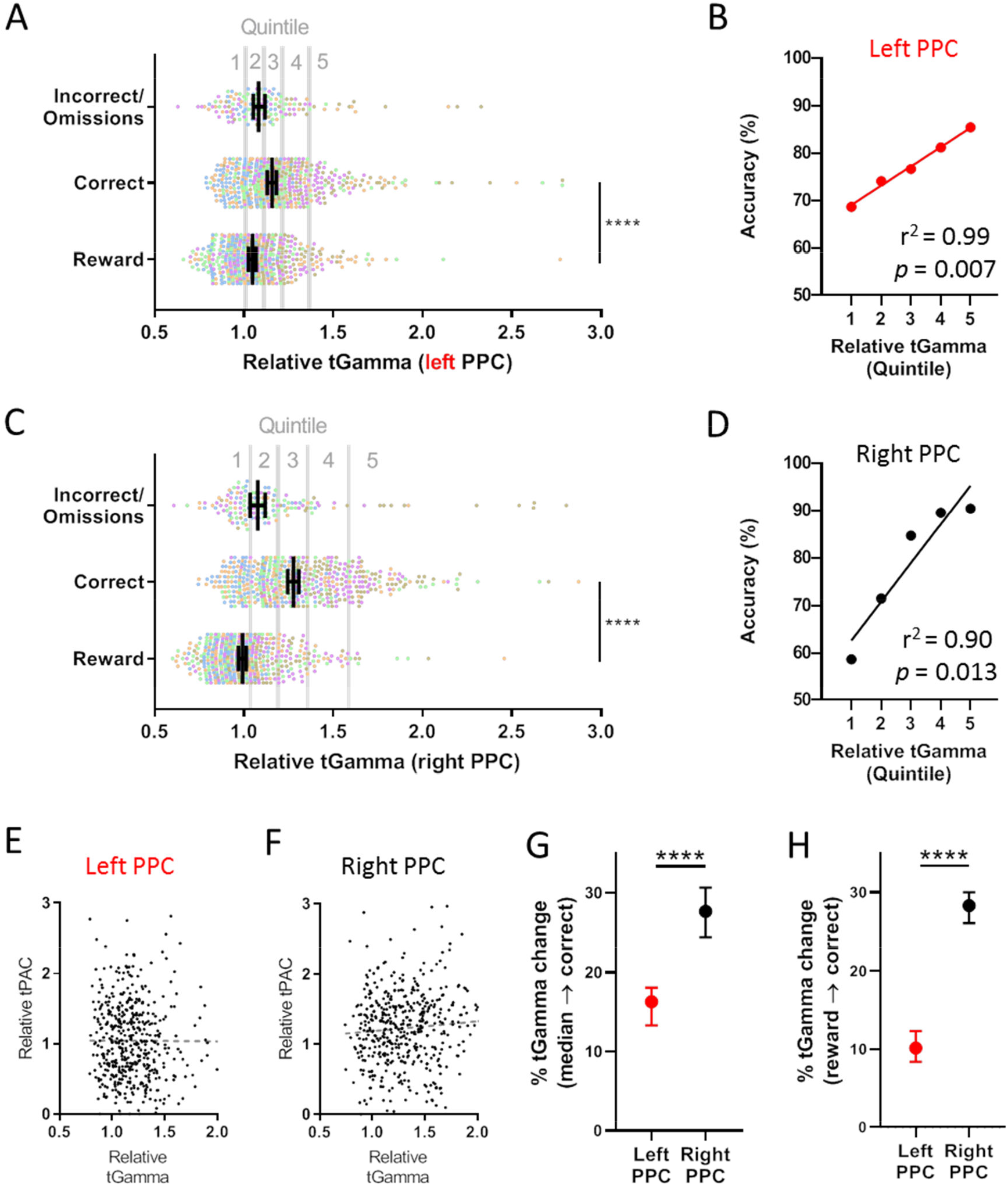
Bilateral positive correlation between posterior parietal tGamma and performance with a difficult task. (A) In the left PPC, there was a significant increase in tGamma prior to correct responses compared to the reward period (n=5 mice, 2 sessions/mouse, \*\*\*\**p*<0.0001). (B) In contrast to the simple task, there was a significant positive correlation between accuracy and tGamma in the left PPC. (C-D) In the right PPC, there was a significant increase in tGamma between the reward period and the ITI prior to a correct response, \*\*\*\**p*<0.0001), (E) No significant relationship between tGamma and tPAC prior to a correct response in the left PPC (r^2^<0.0001, *p*=0.920), and (F) small but significant correlation between tGamma and tPAC in the right PPC (r^2^=0.009, *p*=0.040). The change in tGamma in the right PPC was significantly greater than the left PPC when compared to both (G) median tGamma and (H) tGamma during the reward period.

### No correlation between theta power and accuracy during the attentive state

To dissect the role of theta power further, we next analyzed time-resolved theta (tTheta) with respect to performance in the 5-CSRTT (**Supplemental Figure 3**). In the simple task (SD1, fixed ITI), tTheta increased significantly from the reward period to the ITI prior to both correct and incorrect/omission responses, but there was no correlation with accuracy in either hemisphere (**Supplemental Figure 3A-D**). There remained a right hemispheric dominance in the increase in theta power prior to a correct response when compared to either median tTheta power or the reward period (**Supplemental Figure 3E-F**). With both left and right PPC tTheta, there was a significant increase between the reward and correct periods (*p*<0.0001, nested one-way, **Supplemental Figure 3G, I**). Despite these increases in theta power, there was no significant relationship between tTheta and accuracy in either left or right PPC (**Supplemental Figure 3H, J**). When directly comparing left and right PPC tTheta in a difficult task, right PPC tTheta was significantly greater than left PPC tTheta prior to a correct response when compared to the reward period (p*<*0.0001) but not to median tTheta levels (*p*=0.340, **Supplemental Figure 3K, L**). In addition, tTheta had no significant correlation with tPAC in both right and left PPC for both simple and difficult tasks (r^2^<0.01, *p*>0.05). These findings indicate that theta power increased to a greater degree in the right PPC than the left PPC prior to a correct response, but elevated theta power alone was not sufficient to improve performance on a task of sustained attention.

### Unique changes in tTheta, tGamma and tPAC in a difficult task compared to a simple task

We next specifically assessed whether task difficulty influenced changes in rhythmic dynamics prior to a correct response during the 5-CSRTT. Compared to the reward period, there was a significantly greater increase in tTheta prior to a correct response in the difficult task when compared to the simple task in both left and right PPC (Left PPC: 32.7% [24.5-36.7, 95% CI] with SD1 and 54.9% [43.8-70.2, 95% CI] with SD0.6, *p*=0.008; Right PPC: 43.1% [36.5-47.2, 95% CI] with SD1 and 72.8% [61.3-81.2, 95% CI] with SD0.6; *p*=0.001, nested one-way ANOVA with Sidak’s test for multiple comparisons, **Figure 7A**). tGamma was also increased in both left and right PPC in SD0.6 compared to SD1 but not to a significant degree (Left PPC: 7.3% [1.4-14.0, 95% CI] with SD1 and 10.8% [9.1-13.9, 95% CI] with SD0.6; Right PPC: 23.8% [20.0-27.1, 95% CI] with SD1 and 30.0% [27.6-31.8, 95% CI] with SD0.6; **Figure 7A**). In contrast, there was a trend toward reduced tPAC in the left PPC and elevated tPAC in the difficult task compared to the simple task, suggesting a widening hemispheric asymmetry favoring the right PPC (Left PPC: 18.9% [10.3-24.4, 95% CI] with SD1 and 15.2% [2.9 to 19.6, 95% CI] with SD0.6; Right PPC: 31.3% [27.1-37.4, 95% CI] with SD1 and 47.2% [36.1-53.9, 95% CI] with SD0.6; **Figure 7A**). To more directly assess differences in hemispheric asymmetry, an asymmetry index for tTheta, tGamma, and tPAC was created by taking the difference between the relative magnitude of change in left and right PPC, with a more negative result indicating a greater rightward bias (see Methods). Interestingly, while there was no significant change in the asymmetry index for tTheta or tGamma between the simple and difficult tasks, hemispheric asymmetry for tPAC favoring the right PPC became significantly greater in SD0.6 (−0.11 [−0.18 to −0.07, 95% CI] with SD1 and −0.37 [−0.50 to −0.21, 95% CI], *p*=0.019, nested one-way ANOVA with Sidak’s correction for multiple comparisons, **Figure 7B**). Altogether, these findings indicate that a greater task difficulty is associated with a symmetric, bilateral increase in theta power, but an asymmetric increase in TG-PAC, dominated by the right PPC.

**Figure 7.**
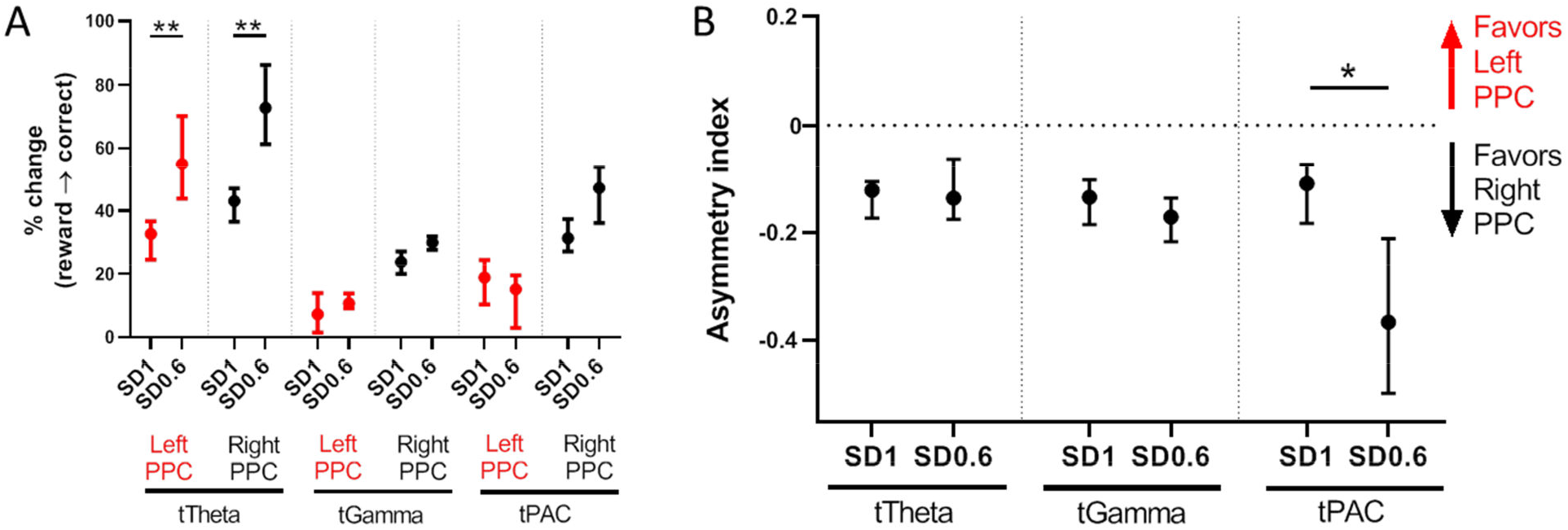
Significantly increased hemispheric asymmetry favoring TG-PAC in the right PPC with greater task difficulty. (A) Increased tTheta in both left and right PPC in the difficult (SD0.6) compared to the simple (SD1) task (\*\**p*<0.01, nested one-way ANOVA with Sidak’s test for multiple comparisons). (B) Hemispheric symmetry with tTheta and tGamma remains unchanged despite increased task difficulty; however, tPAC becomes significantly more asymmetric in the difficult task, favoring the right PPC (\**p*=0.019).

## Discussion

In this study, we report on the dynamic changes of theta power, gamma power and theta-gamma phase-amplitude coupling (TG-PAC) in the PPC in mice during a sustained attention task. We found that (i) theta-gamma phase-amplitude coupling (TG-PAC) was consistently stronger in the right compared to left PPC, regardless of task difficulty, (ii) during the highly attentive state (the inter-trial interval prior to a correct response), both gamma power and TG-PAC were independently significantly elevated in the right PPC and consistently correlated with attention performance; and (iii) with increased task difficulty – theta power symmetrically increased bilaterally, gamma power in bilateral PPC correlated with attention performance, and hemispheric asymmetry in TG-PAC became significantly more pronounced, favoring the right PPC.

### Lateralization of sustained attention to the right posterior parietal cortex

We found a strong dominance of right (compared to left) PPC gamma power and TG-PAC during a sustained attention task. Lateralization of sustained attention toward the right PPC has been supported by great deal of evidence from lesion and functional imaging studies in humans (Petersen and Posner, 2012; Posner and Petersen, 1990); however, the dominance of the right PPC for sustained attention in rodents, to our knowledge, has not previously been recognized. Our findings are consistent with prior studies showing lateralization of other higher-order cortical function in mice as seen in humans (Duboc et al., 2015), including right hippocampal dominance for visuospatial memory (Shinohara et al., 2012) and left hemispheric dominance for vocalization expression and response (Doran et al., 2015; Ehret, 1987; Levy et al., 2019). Altogether, our results join a growing body of data positioning mice as a potential translational model of some higher order cognitive functions.

Human neocortical theta (4-7 Hz) is an overlapping but overall lower frequency range than mouse theta (6-10 Hz), and the neocortical theta rhythm in rodents can largely reflect volume conduction from hippocampal theta (Buzsáki, 2002; Cantero et al., 2003; Nishida et al., 2004; Watrous et al., 2013). With these caveats, in patients with epilepsy who were implanted with intracranial EEG, Szczepanski et al have previously shown a high degree of delta/theta (2-5 Hz) to high gamma (80-250 Hz) coupling in frontoparietal regions during a sustained attention task (Szczepanski et al., 2014). A trend toward greater PAC in the right hemisphere compared to the left hemisphere was found, but statistical significance may have been limited by aggregating all sites within the frontoparietal network and a lack of comparison between hemispheres in the same patient. Further study of ECoG in patients with bilateral sampling from parietal regions is warranted to determine if PAC is consistently lateralized to the right hemisphere in humans during an attention task.

### Positive correlation of right posterior parietal TG-PAC and gamma power with attention performance

Right PPC TG-PAC was positively correlated with performance in mice with both a simple and difficult attention task. Accumulating evidence points to PAC as an efficient computational mechanism for integrating top-down and bottom-up pathways in the cortex (Canolty and Knight, 2010; Wang et al., 2014). In the limbic system, PAC in the rat hippocampus has a positive correlation with learning behavior (Tort et al., 2009); PAC in rat orbitofrontal cortex has a positive correlation with olfactory decision making (van Wingerden et al., 2014); and PAC in the basolateral amygdala has a positive correlation with freezing behavior (Stujenske et al., 2014). Another recent study found a positive correlation between prefrontal cortex (but not visual cortex) TG-PAC and sustained attention using a Go-No Go task (Han et al., 2019); however, compared to the present study, the frontal electrodes were more anterolaterally placed, and data from the hemispheres were averaged, limiting the ability to discriminate hemispheric dominance. Nonetheless, there is a consistent theme of PAC correlating with a behavior that is specific to the function of its associated cortical region in rodents.

Gamma power also showed a right-sided dominance, significantly increasing prior to a correct response to a greater degree in the right PPC than in the left PPC. Gamma oscillations have been associated with enhanced sensory processing (Saalmann et al., 2007). Importantly, only the increase in gamma power in the right PPC correlated significantly with attention performance with a simple task. With a difficult task, however, the increase in gamma power correlated significantly with accuracy in both left and right PPC. These findings suggest that more difficult or complex tasks may recruit the left hemisphere to assist in sustaining attention. Similarly, within the auditory cortex, the right hemisphere is activated by more generic tones, but the left hemisphere becomes activated by more complex vocalizations (Levy et al., 2019).

The enhanced correlation between left PPC gamma power and accuracy with a difficult task is hard to reconcile with the simultaneous increase in hemispheric asymmetry of TG-PAC favoring the right hemisphere. However, given the lack of significant correlation between gamma power and TG-PAC in both simple and difficult tasks, gamma power and TG-PAC in the left PPC need not change in the same direction. Despite the positive correlation between left PPC gamma power and accuracy, right PPC gamma power still increased to a significantly greater degree than left PPC gamma power, thus showing continued right PPC dominance during the difficult task.

### Importance of theta phase but not amplitude in attention performance

Given the near-zero correlation between TG-PAC and gamma power during the attentive phase (inter-trial interval) prior to a correct response, the association of TG-PAC with accuracy cannot be explained by increased gamma power alone. The coordination of network activities between distant sites occurs in the theta frequency range both during the resting state and the attentive state (Marek and Dosenbach, 2018), and theta rhythms have specifically been shown to aid in rhythmic sampling of the environment with large scale communication between frontal and parietal regions during sustained attention tasks (Fiebelkorn and Kastner, 2019; Sellers et al., 2016). In this study, the power of theta significantly increased bilaterally in a difficulty-dependent manner during the attentive period of the task, similar to prior work in ferrets (Sellers et al., 2016), but changes in theta power had no significant correlation with accuracy.

There are some limitations to this study. First, the results from these experiments show only a correlation between posterior parietal TG-PAC, gamma power, and sustained attention. To establish a causal link, further work should aim to directly alter posterior-parietal theta-gamma phase-amplitude coupling using pharmacological, electrical, optogenetic, and/or pharmacogenetic interventions while evaluating changes in sustained attention. Second, only male mice were used; future studies should incorporate both male and female subjects to increase the generalizability of our findings. Third, since the mice progressed in a linear fashion through difficulty, some of the effects may have been due to practice; replication studies should randomize simple and difficult task order after performance has stabilized to account for these potential time-varying factors. Finally, the main focus of this work has been on the posterior parietal cortex. While there were electrodes placed superficially over Supplemental motor cortex, there was no clear modulation of TG-PAC with sustained attention in either hemisphere. Further evaluation in bilateral medial and lateral prefrontal cortices may help to delineate to what degree hemispheric dominance may be present throughout the frontoparietal network in mice.

Our results in mice are consistent with convergent evidence across species of a right hemispheric dominance for sustained attention. These findings implicate the potential for modulation of theta-gamma coupling and gamma power in the right PPC as a means to modulate attention performance. Future investigation may probe the effect of pharmacology, neuromodulation, and neurofeedback on these electrophysiologic signatures to potentially aid in disorders of attention.

---

**Supplemental Figure 1.**
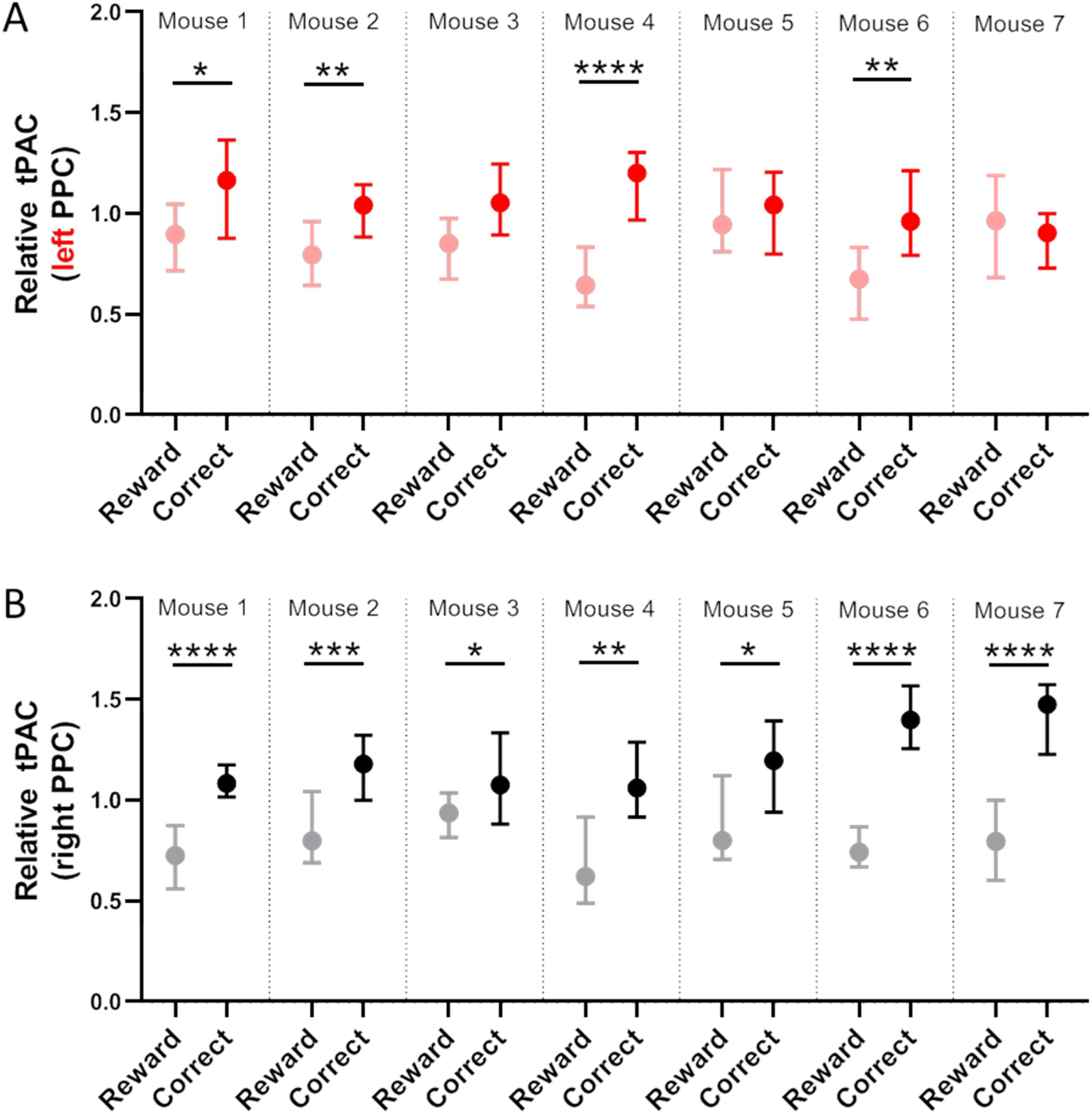
Changes in relative tPAC in individual mice at SD1. (A) In left PPC, there was a significant increase in relative tPAC from the reward period to the intertrial interval prior to correct responses in 4 out of 7 mice. (B) In right PPC, all 7 mice had a significant increase in relative tPAC prior to a correct response (\**p*<0.05, \*\**p*<0.01, \*\*\**p*<0.001, \*\*\*\**p*<0.0001, Wilcoxon matched-pairs signed rank test).

**Supplementary Figure 2.**
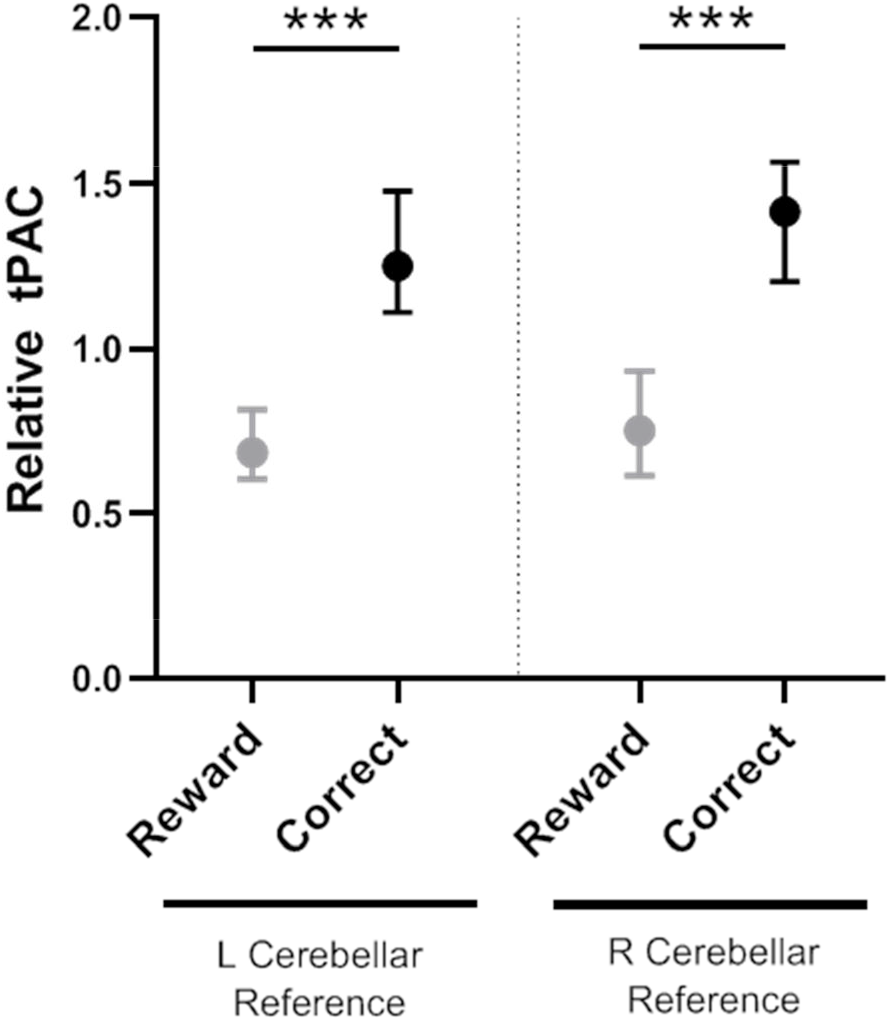
Significant increase in relative tPAC independent of referenced cerebellar hemisphere. In one mouse, left and then right cerebellar references were used on consecutive days, and the mouse passed criteria for SD1 in both sessions. tPAC in the right PPC significantly increased to a similar degree despite the change in reference (\**p*<0.001, Wilcoxon matched-pairs signed rank test).

**Supplementary Figure 3.**
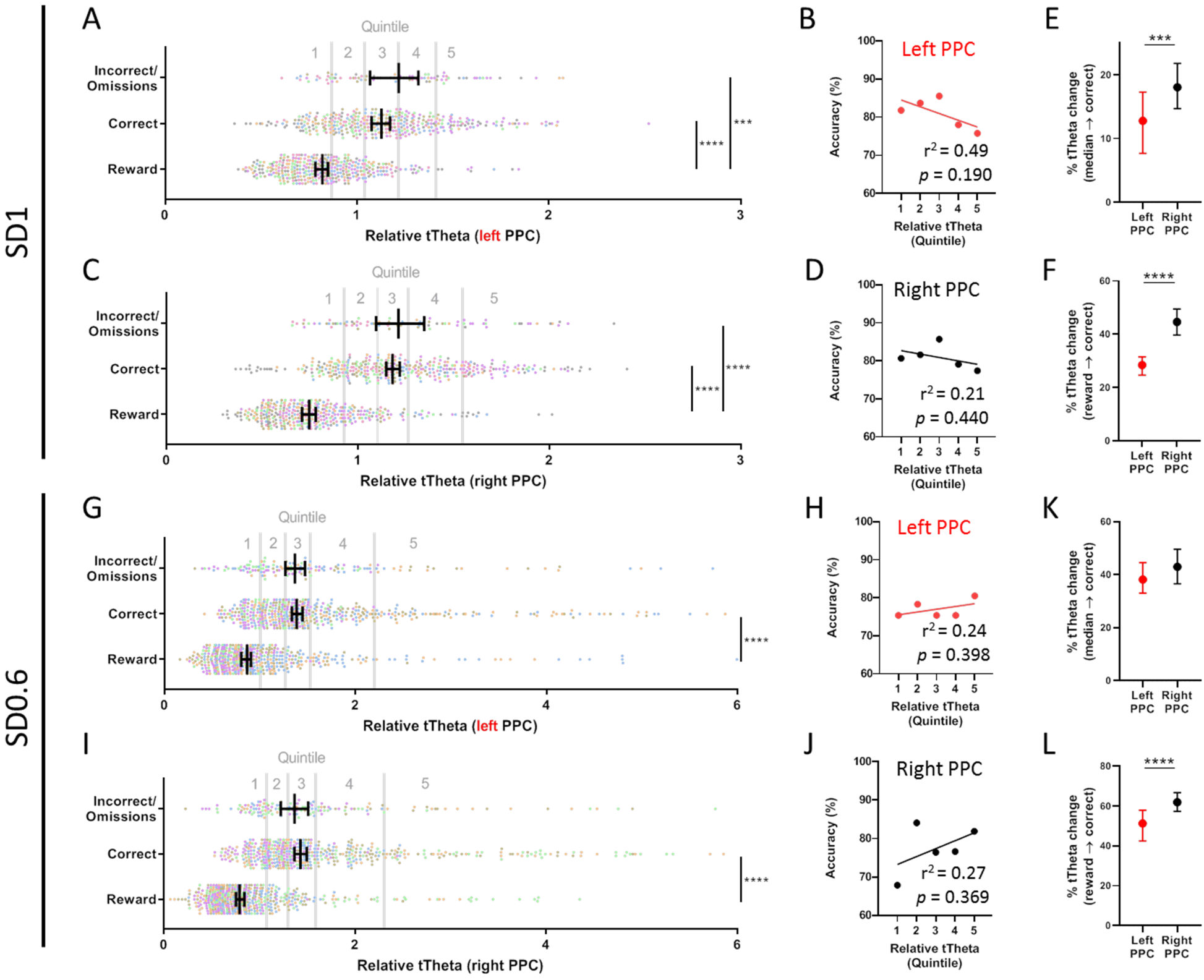
No relationship between theta power and accuracy in both simple and difficult tasks. (A-D) In both left and right PPC during SD1 (simple task), tTheta increased significantly from the reward period to the ITI before both correct and incorrect/omission responses (\*\*\**p*<0.001, \*\*\*\**p*<0.0001); however, there was no significant relationship between tTheta and accuracy. (E, F) Significantly greater theta power prior to correct responses when compared to either median theta power or theta power during the reward period. (G-J) In both left and right PPC in SD0.6 (difficult task), there was a significant increase in theta power between the reward period and the ITI prior to correct responses, but no significant correlation between accuracy and tTheta (\*\*\**p*<0.0001). (K) No significant difference in the change in tTheta between the left and right PPC when correct responses are compared to median tTheta; (L) however, there was a significant increase in tTheta prior to a correct response when compared to the ITI prior to reward responses (\*\*\*\**p*<0.0001).

